# Metabolic Reprogramming in Rheumatoid Arthritis Synovial Fibroblasts: a Hybrid Modeling Approach

**DOI:** 10.1101/2022.07.20.500752

**Authors:** Sahar Aghakhani, Sylvain Soliman, Anna Niarakis

## Abstract

Rheumatoid Arthritis (RA) is an autoimmune disease characterized by a highly invasive pannus formation consisting mainly of synovial fibroblasts (RASFs). This pannus leads to cartilage, bone and soft tissue destruction in the affected joint. RASFs’ activation is associated with metabolic alterations resulting from dysregulation of extracellular signals transduction and gene regulation machinery. Deciphering the intricate mechanisms at the origin of this metabolic reprogramming may provide significant insight into RASFs’ involvement in RA’s pathogenesis and offer new therapeutic strategies. Qualitative and quantitative dynamic modeling can address some of these features, but hybrid models represent a real asset in their ability to span multiple layers of biological machinery. This work presents the first hybrid RASF model: the combination of a cell-specific qualitative regulatory network with a global metabolic network. The automated framework for hybrid modeling exploits the regulatory network’s trap-spaces as additional constraints on the metabolic networks. Subsequent flux balance analysis allows assessment of RASFs’ regulatory outcomes’ impact on their metabolic flux distribution. The hybrid RASF model simulates the experimentally observed metabolic reprogramming induced by signaling and gene regulation in RASFs. Simulations also enable further hypotheses on the potential reverse Warburg effect in RA. RASFs may undergo metabolic reprogramming to turn into “metabolic factories”, producing high levels of energy-rich fuels and nutrients for neighboring demanding cells through the crucial role of HIF1.

**Author Summary:** We successfully built the first large-scale hybrid dynamical model for human Rheumatoid Arthritis Synovial Fibroblasts (RASFs) including signaling, gene regulation and metabolism. We used a state-of-the-art molecular map for upstream signaling and gene regulation, the tool CaSQ to infer a large-scale Boolean model, and a genome-scale metabolic model. Trap-spaces of the Boolean asynchronous model were used to infer additional metabolic constraints on the metabolic network for subsequent flux balance analysis. This method allowed us to study the impact of various regulatory initial conditions on RASFs’ metabolic fluxes distribution. Our model successfully reproduces the metabolic reprogramming of RASFs which shift their ATP production from oxidative pathways to glycolysis, highlighting the key role of HIF1 in this process. Our findings allow us to hypothesize a reverse Warburg relationship occurring between RASFs and other RA joint cells. Similarly to tumor microenvironment’s fibroblasts, RASFs would undergo a metabolic switch and reprogram their metabolism to adapt to their hypoxic environment and provide crucial metabolic intermediates to neighboring cells to sustain their inflammatory activity.

## 1. Introduction

Rheumatoid Arthritis (RA) is a chronic auto-immune disorder affecting approximately 0.5-1% of industrialized countries’ population (Scott et al., 2010). The growing proportion of people affected (Safiri et al., 2019) and the impact on the patients’ quality of life have turned RA into a severe public health issue. Resulting from the progressive destruction of cartilage and bones, RA leads to pain, swelling and stiffness in the affected joint (Scott et al., 2010). In addition, extra-articular manifestations often follow, such as cardiac, renal or neurological disorders, all associated with excess mortality (Figus et al., 2021).

Although it is widely acknowledged that an immune system’s dysfunction triggers RA, the etiology of the disease is not yet fully elucidated. RA is a complex disease resulting from multiple intertwined factors (McInnes & Schett, 2011). Genetic factors of susceptibility were initially identified (Yarwood et al., 2014), along with epigenetic factors, namely DNA methylation or histone proteins modifications (Nemtsova et al., 2019). Environmental factors have also been recognized recently, including pollution (Sigaux et al., 2019) and smoking (Chang et al., 2014). Finally, RA’s female-to-male prevalence ratio having consistently been established at 3:1 (Lawrence, 1970), hormonal factors are suspected to play a role (Alpízar-Rodríguez et al., 2016). This complexity and the fragmented knowledge of the disease pathophysiology contribute to the lack of a cure for RA. The different treatment options aim to reduce further damage and symptoms to improve the patient’s overall quality of life. Disease-modifying antirheumatic drugs (DMARDs) including hydroxychloroquine or methotrexate, may be used to promote remission by slowing or stopping the progression of the disease (Smolen et al., 2005). Nonsteroidal anti-inflammatory drugs (NSAIDs) such as acetylsalicylate, naproxen, and ibuprofen may also be considered to relieve pain and decrease inflammation (Ong et al., 2007). A new lead lies in the action of JAK inhibitors, namely tofacitinib, to remedy inflammation (Tanaka & Yamaoka, 2013b). However, despite the various options available, a significant proportion of RA patients do not respond significantly to treatment and are in a state of therapeutic distress (J. S. Smolen et al., 2018).

RA’s pathogenesis is associated with synovial hyperplasia or pannus formation, consisting of the accumulation of macrophages and synovial fibroblasts (RASFs), leading to inflammation and bone erosion (Müller-Ladner et al., 2007). RASFs are the most common resident cells of the synovial membrane and play a crucial role in the onset and disease progression (Bottini & Firestein, 2012). They are derived from the joints’ synovial membrane and differ from healthy fibroblasts’ morphology, gene expression pattern and phenotype. In a healthy joint, fibroblasts are arranged as a one or two-cell layer and are responsible for the synovial membrane’s structural integrity (Castor, 1960). In addition, fibroblasts ensure nutrient supply and joint lubrication and create the synovial fluid’s non-rigid extracellular matrix, aiding wound healing and tissue repair. On the contrary, the rheumatic pannus is a 10-15 cell depth layer (Kiener et al., 2010) where RASFs present an aggressive tumor-like phenotype. They express very high levels of cytokines, chemokines and matrix-degrading enzymes, causing cartilage damage and maintaining inflammation. Their reduced contact inhibition and their resistance to apoptosis allow them to grow massively, invading periarticular tissues and contributing to their destruction. In addition, RASFs are considered as primary drivers of angiogenesis in RA joints (Bottini & Firestein, 2012; Turner & Filer, 2015). In light of these advances, RASFs, once considered passive bystanders, are now recognized as active players in disease pathogenesis. Therefore, RASF-directed therapies could become a complementary approach to currently used immune therapies (Filer, 2013).

Healthy fibroblasts’ transformation into RASFs appears to be associated with metabolic alterations (Aghakhani et al., 2020). Under normoxic conditions, healthy cells oxidize one glucose molecule into two pyruvate molecules along with 36 molecules of adenosine triphosphate (ATP) through glycolysis followed by tricarboxylic acid cycle (TCA) and oxidative phosphorylation (OXPHOS). In hypoxic conditions, pyruvate is diverted from TCA and transformed into lactate, generating two molecules of ATP. In cancer cells, this metabolic switch can also occur in the presence of oxygen, known as the Warburg effect (Warburg et al., 1927). Cancer cells opt for glycolysis to produce energy in the form of ATP even if it is less efficient because glycolytic intermediates will feed metabolic pathways supporting cell growth, proliferation and survival (e.g., pentose phosphate pathway (PPP), fatty acids, glutaminolysis). A few recent experimental studies indicated a link between altered metabolism and inflammation levels in RA. Measurements of synovial tissue PO2 levels demonstrated that the RA joint microenvironment is profoundly hypoxic (Ng et al., 2010). In addition, increased glucose uptake, glycolytic enzymes, lactate secretion, and oxidative damage have been identified in RASFs, correlating with cytokine levels and disease scores (Fearon et al., 2018).

While provided with some understanding of the metabolic pathways altered in RASF, there is still a lack of insight into their activating stimuli and the relevance of such metabolic reprogramming in the RA joint. As fragmented as it may be, a collection of the knowledge in terms of extra- and intra-cellular signaling along with gene regulation pathways involved in RASFs’ metabolic reprogramming would be beneficial. Efforts have already been made in this direction with the publication of the first RA-map (Singh et al., 2020), followed by the RA-Atlas (Zerrouk et al., 2022). The RA-Atlas is an interactive, manually curated representation of molecular mechanisms involved in RA’s pathogenesis following Systems Biology Graphical Notation (SBGN) standards (Le Novère et al., 2009). It includes an updated version of the RA-map, the RA-map V2, with the addition of metabolic machinery, and cell-specific molecular interaction maps for CD4+ Th1 cells, fibroblasts, M1 and M2 macrophages. Although including the impact of signaling and gene regulation on four major metabolic pathways, namely glycolysis, PPP, TCA and OXPHOS, the RA-map V2 is limited by its non-cellular specificity and static knowledge base function. However, it can be used to construct dynamic computational models to extend knowledge with executable information.

Dynamic modeling is necessary to understand the emergent behavior of biological entities when complex and intertwined pathways are involved. It helps elucidate complex mechanisms occurring at different scales (e.g., signaling, gene regulation, and metabolism) between different entities (e.g., ligands, receptors, proteins, metabolites, genes, RNAs). While dynamic models based on quantitative information provide great insight into biological systems, qualitative models based on logical relationships among components provide an appropriate description for systems with unknown mechanistic foundations or lacking precise quantitative data (Saadatpour & Albert, 2016). Qualitative Boolean models allow the parameter-free study of the underlying dynamic properties of large-size biological pathways. In Boolean formalism, nodes represent regulatory components, and arcs represent their interactions. Each regulatory component is associated with a Boolean value (0 or 1), indicating its qualitative concentration (absent or present) or its activity level (inactive or active). The state of each node depends on the state of its upstream regulators and is described by a Boolean rule defined by the logical operators “AND”, “OR” and “NOT”. Despite being well-suited to account for the modeling of signaling and gene regulation mechanisms, qualitative Boolean modeling is not appropriate to assess quantitative metabolic properties. Where signaling and gene regulation carry signal flow, metabolism generates mass flow. A widely used method for analyzing metabolic networks is Flux Balance Analysis (FBA). FBA is a mathematical method used in large-scale reconstructions of metabolic networks (Orth et al., 2011). Its main advantage lies in the need for little information regarding enzymes’ kinetic parameters and metabolites’ concentrations as it calculates the flow of metabolites by assuming steady-state conditions.

As outlined above, the various biological features of signaling, gene regulation, and metabolism are usually studied separately through different modeling formalisms. However, considering that a biological phenotype results from their interoperation, a more hybrid formalism capable of spanning these layers would be highly beneficial in studying complex diseases (Gonçalves et al., 2013; Stéphanou & Volpert, 2015; Cruz & Kemp, 2021). One of the recent efforts in this direction is the FlexFlux framework (Marmiesse et al., 2015) combining FBA and qualitative simulations by seeking regulatory steady-states through synchronous updates of multi-state qualitative initial values. Van der Zee & Barberis, 2019 also proposed a framework to integrate Boolean with constraint-based models of metabolism.

Hybrid modeling has already been suggested in RA to improve understanding of the disease, notably with a framework describing pannus formation (Macfarlane et al., 2021), but, to our knowledge, never in a cell-specific manner. In this regard, this work presents the first hybrid RASF model. It was obtained by combining a cell-specific asynchronous Boolean network covering gene regulation and signaling machinery with a global constraint-based metabolic network. This hybrid model allows bridging the gap between various biological features to assess the impact of RASFs’ regulatory outcomes on its central metabolic flux distribution.

## 2. Methods

### 2.1. RASF regulatory model

#### 2.1.1. Inference of a Boolean model from the RA-map V2

The RA-map V2, included in the RA-Atlas (Zerrouk et al., 2022), provided the starting point for this work. It illustrates the central signaling, gene regulation and metabolic pathways (i.e., glycolysis, PPP, TCA and OXPHOS), along with molecular mechanisms and phenotypes involved in RA’s pathogenesis. The RA-map V2 is a global map, gathering information from several cell types, tissues and fluids such as RASFs, synovial tissue, synovial fluid, blood and serum components, peripheral blood mononuclear cells, chondrocytes and macrophages. CaSQ (Aghamiri et al., 2020) was used to infer a Boolean model from the RA-map V2 in the standard Systems Biology Marked up Language-qualitative (SBML-*qual*) (Chaouiya et al., 2013) format. Its latest functionality enables extraction and translation into a model, not an entire molecular map, but only a subpart of interest. In this case, the focus was on extracting pathways upstream and downstream of RASF-specific extracellular ligands, metabolites, and microRNA experimentally demonstrated to be significant in RA’s pathogenesis.

#### 2.1.2. Cellular-specificity assessment

The cellular-specificity of the model was evaluated by comparing its components’ cellular origin to eight different sample-type specific lists provided in the RA-Atlas publication (Zerrouk et al., 2022). They are specific to RASFs, synovial tissue, synovial fluid, blood and serum components, peripheral blood mononuclear cells, chondrocytes, and macrophages.

#### 2.1.3. Annotation score calculation

Thorough bibliographic annotations of the RA-map V2 in the form of PubMed IDs (PMIDs) following MIRIAM (Minimum Information Required In The Annotation of Models) standards (Le Novère et al., 2005), were kept in the associated model. They allowed to calculate annotation scores for each compound present in the model based on the number of bibliographic references describing it.

#### 2.1.4. Regulatory model’s behavior validation

The model’s behavior was assessed to confirm its biological relevance. Simulations were performed on CellCollective (Helikar et al., 2012), an interactive platform to simulate and analyze biological models. Experimental evidence for RASF-specific scenarios was retrieved from *in-vitro* and *in-vivo* studies, in humans when possible, but mostly in murine models of RA. Experimental conditions were used as the model’s inputs, and its outputs were compared to the biological outcome to confirm or refute a scenario’s validation. Two approaches were carried out to assess the model’s behavior. First, a generic validation was performed to verify specific compounds’ contribution to signaling or gene regulation pathways. This mechanistic verification was performed in a synthetic state of the model where all nodes’ values were set to 0 (i.e., absent/inactive), and specific compounds to test were set alternatively to 0 and 1. Secondly, the global behavior of the model was analyzed under RASF-specific initial conditions identified from the literature. This global analysis was conducted at the level of the model’s phenotypes. It allowed to evaluate the different pathways’ interconnection and compare the global model’s behavior to RASFs’ experimentally expected behavior. In both cases, simulations were performed in the asynchronous updating mode, with a simulation speed of one and a sliding window of 30.

### 2.2. Metabolic model

The metabolic network used in this work is the MitoCore model (Smith et al., 2017), a manually curated constraint-based model of human metabolism. This genome-scale metabolic model includes two compartments (i.e., cytosol and mitochondria), 74 metabolites, 324 metabolic reactions, and 83 transport reactions. The default parameters of MitoCore simulate normal cardiomyocyte metabolism. However, the default simulation settings can be applied to various biological contexts without necessarily implying cell-specific features. Indeed, cardiomyocytes can metabolize a wide range of substrates, have reactions common to many other cell types, and represent the metabolism of the human heart, an organ of utmost importance in human health, disease and toxicology. Moreover, MitoCore includes processes that are inactive in cardiomyocytes but that can be activated to represent other cell-type’s metabolic features (e.g., gluconeogenesis, ketogenesis, β-alanine synthesis and folate degradation). As a proof of concept regarding the generalization of their model, MitoCore modelers were able to successfully simulate the fumarase deficiency, a nervous system condition, using the default cardiomyocyte parameters.

### 2.3. Framework for hybrid modeling

The general architecture of the framework for hybrid modeling is provided in Figure 1.

**Figure 1.**
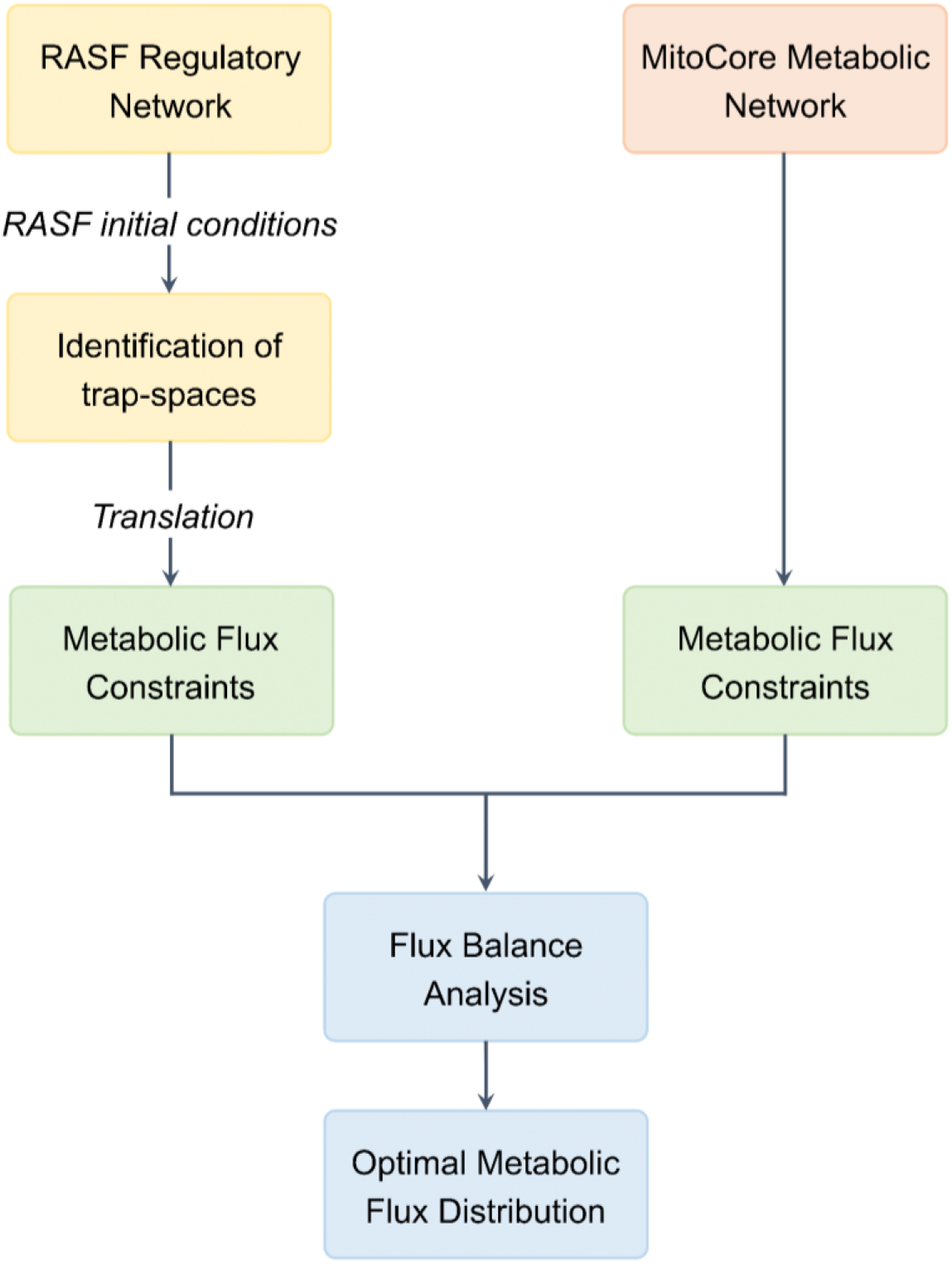
General architecture of the hybrid modeling framework.

#### 2.3.1. Value propagation

Value propagation (Saadatpour et al., 2013; Hernandez et al., 2020), a method implemented in the CoLoMoTo notebook (Naldi et al., 2018), was applied to the qualitative regulatory model to facilitate its analysis. When given a set of logical rules and a cellular context, this iterative algorithm allows the computation of specific components’ dynamical consequences on the model’s behavior. It reveals the influence specific compounds may exert on the network’s dynamics. Note that this method does not impact the asymptotic behavior of the model, all dynamical consequences calculated in this manner would occur regardless.

#### 2.3.2. Identification of regulatory trap-spaces

Evaluating the influence of RASF-specific components on the global model’s behavior through value propagation allows decreasing its complexity to identify trap-spaces. Trap-spaces, also called stable motifs or steady symbolic states, are parts of the dynamics from which the system cannot escape (Zañudo & Albert, 2013; Klarner et al., 2014). A trap-space is a partially assigned state such that all possible successors of all states which belong to the motif also belong to the motif. Thus, minimal trap-spaces (later referred to as “trap-spaces” only for readability) offer a good approximation of complex attractors and faithfully capture the asymptotic behavior of Boolean models.

The outputs of value propagation were considered as a new set of initial conditions and the `biolqm.trapspace` function was used to identify the model’s trap-spaces. This method implements a constraint-solving method based on decision diagrams for efficiently identifying a model’s trap-spaces without performing simulation.

#### 2.3.3. Projection of metabolic regulatory trap-spaces

The regulatory model does not allow to differentiate a protein with a signaling function from a metabolic enzyme or any simple molecule from a metabolite. To overcome this limitation, all regulatory model’s components were extracted as a list, as well as MitoCore’s enzymes, and MitoCore’s metabolites. Both regulatory and metabolic network’s metabolic components being consistently named with BiGG IDs (King et al., 2015), the regulatory model’s components were compared to MitoCore’s metabolites and a list of the regulatory model’s metabolites was obtained. This matching was limited by excluding a list of manually predefined compounds considered by MitoCore as metabolites but actually are common metabolic intermediates (Table S1, appendix). Similarly, a list of the regulatory model’s enzymes was extracted. Once obtained, those lists allowed to project previously identified regulatory trap-spaces on the metabolic enzymes and metabolites.

#### 2.3.4. Constraining metabolic fluxes

The maximum value of trap-spaces relative to metabolic compounds allows constraining associated metabolic flux. A metabolic enzyme-associated maximal trap-space value strictly superior to 0 means this metabolic enzyme might be activated or present according to the regulatory model’s outcome. An enzyme being qualitatively activated or present does not give information about the feasibility nor the kinetics of the metabolic reactions it catalyzes. Similarly, a metabolite-relative maximal trap-space value strictly superior to 0 shows the metabolite is produced in some of these regulatory conditions. However, it does not give information about its producing reactions’ high or low flux. Thus, it is impossible to influence the reaction flux of metabolic compounds associated with maximal trap-spaces values strictly superior to 0.

On the other hand, a metabolic enzyme with a projected maximal trap-space value equal to 0 expresses the inactivation or absence of the said enzyme by signaling or gene regulation pathways. The catalyzed metabolic reactions will not happen. A metabolite-associated maximal trap-space value of 0 denotes the non-production of the said metabolite. Its producing reactions will not occur. Hence, for every metabolic enzyme or metabolite with projected maximal regulatory trap-spaces values equal to 0, the flux of its associated MitoCore metabolic reaction (i.e., catalyzed reactions or producing reactions) is constrained to 0.

### 2.4. Metabolic network analysis

#### 2.4.1. Flux Balance Analysis

Flux Balance Analysis (FBA) was performed using CobraPy (Ebrahim et al., 2013) to evaluate MitoCore’s metabolic flux distribution. Two FBAs were conducted to highlight a potential change in metabolic fluxes’ distribution under RASF-specific conditions. The first FBA was conducted without additional constraints, reflecting the healthy control state. The second FBA was conducted with additional metabolic flux constraints extracted from the regulatory model’s trap-spaces, reflecting RASFs’ condition. The objective function was set to maximum cellular ATP production to reflect the primary energy production role of central metabolism. It was manually defined as the sum of the three cellular ATP-producing reactions (i.e., the seventh and tenth reactions of glycolysis and OXPHOS’ last step).

The MitoCore metabolic model lacking cellular and tissular specificity, a numerical interpretation of metabolic flux values was not possible. FBA results were only interpreted in terms of metabolic flux distribution. For instance, the ratios of ATP production from glycolytic or OXPHOS reactions relative to total ATP production (represented by the objective function) were calculated. In addition, an analysis of uptake and secretion of carbon fluxes (C-flux) was carried out to indicate the central carbon metabolism. The C-flux of an uptake reaction represents the total cellular carbon influx from this specific uptake reaction. Similarly, the C-flux of a secretion reaction represents the proportion of total cellular carbon efflux coming from this specific secretion reaction. Finally, a comparison of both FBAs’ internal fluxes was conducted, and a difference in metabolic flux was identified from a greater than 2-fold variation.

#### 2.4.2. Regulatory initial condition’s knock-out/knock-in simulations

To identify regulatory pathways potentially responsible for RASF’s metabolic alterations, a knock-out/knock-in strategy of the regulatory network’s initial conditions was conducted. All initial regulatory conditions set to 1 in the RASF-specific configuration were successively set to 0 while the other remained at RASF-specific values. Similarly, all regulatory components’ initial conditions set to 0 in the RASF-specific configuration were successively set to 1 while the other remained at RASF-specific values. Subsequent value propagation, metabolic compounds-projected regulatory trap-spaces’ identification and extraction of metabolic constraints were conducted to perform FBA and evaluate metabolic fluxes’ distribution in these new conditions. To depict the metabolism’s key function of energy production, the proportion of total cellular ATP production from glycolysis and OXPHOS were compared in the various FBAs.

## 3. Results

### 3.1. RASF regulatory model

Translation of the RA-map V2 by focusing on RASF-specific molecular pathways (Table 1) provided a dynamic Boolean model of 359 nodes, including 14 inputs, and 642 interactions (Figure S1, appendix).

**Table 1.** RASF-specific components and the export direction used to infer a Boolean model from the RA-map V2 using CaSQ.

| Export direction | Component | Class | Reference |
| --- | --- | --- | --- |
| Upstream | HIF1 | Protein | Brouwer et al., 2009 |
| Downstream | IL6 | Extracellular ligand | Guerne et al., 1989 |
| Downstream | IL18 | Extracellular ligand | Gracie et al., 1999 |
| Downstream | FGF1 | Extracellular ligand | Qu et al., 1995 |
| Downstream | PDGFA | Extracellular ligand | Rosengren et al., 2010 |
| Downstream | TGFB1 | Extracellular ligand | Cheon et al., 2002 |
| Downstream | WNT5A | Extracellular ligand | Sen et al., 2001 |
| Downstream | SFRP5 | Extracellular ligand | Kwon et al., 2010 |
| Downstream | RANKL | Extracellular ligand | Takayanagi et al., 2000 |
| Downstream | IL17A | Extracellular ligand | Kotake et al., 1999 |
| Downstream | FASLG | Extracellular ligand | Fukuda et al., 2021 |
| Downstream | IKBA/NFKB1/RELA | Extracellular ligand | Nejabatbakhsh Samimi et al., 2020 |
| Downstream | TNF | Extracellular ligand | Lee et al., 2013 |
| Downstream | GLC | Extracellular metabolite | Garcia-Carbonell et al., 2016 |
| Downstream | MIR192 | microRNA | Li et al., 2017b |

Although the RA-map V2 is a collection of information from several cell types, fluids and tissues, the inferred model is mostly RASF-specific (82%) (Figure 2A). When interpreting those results, one must consider that a specific component can be common to several cell types. If only exclusive components are considered, the model is 91% RASF-specific (Figure 2B). This high level of cellular specificity, along with the phenotypes and extracellular ligands specificity, allows referring to this model as the RASF model.

**Figure 2.**
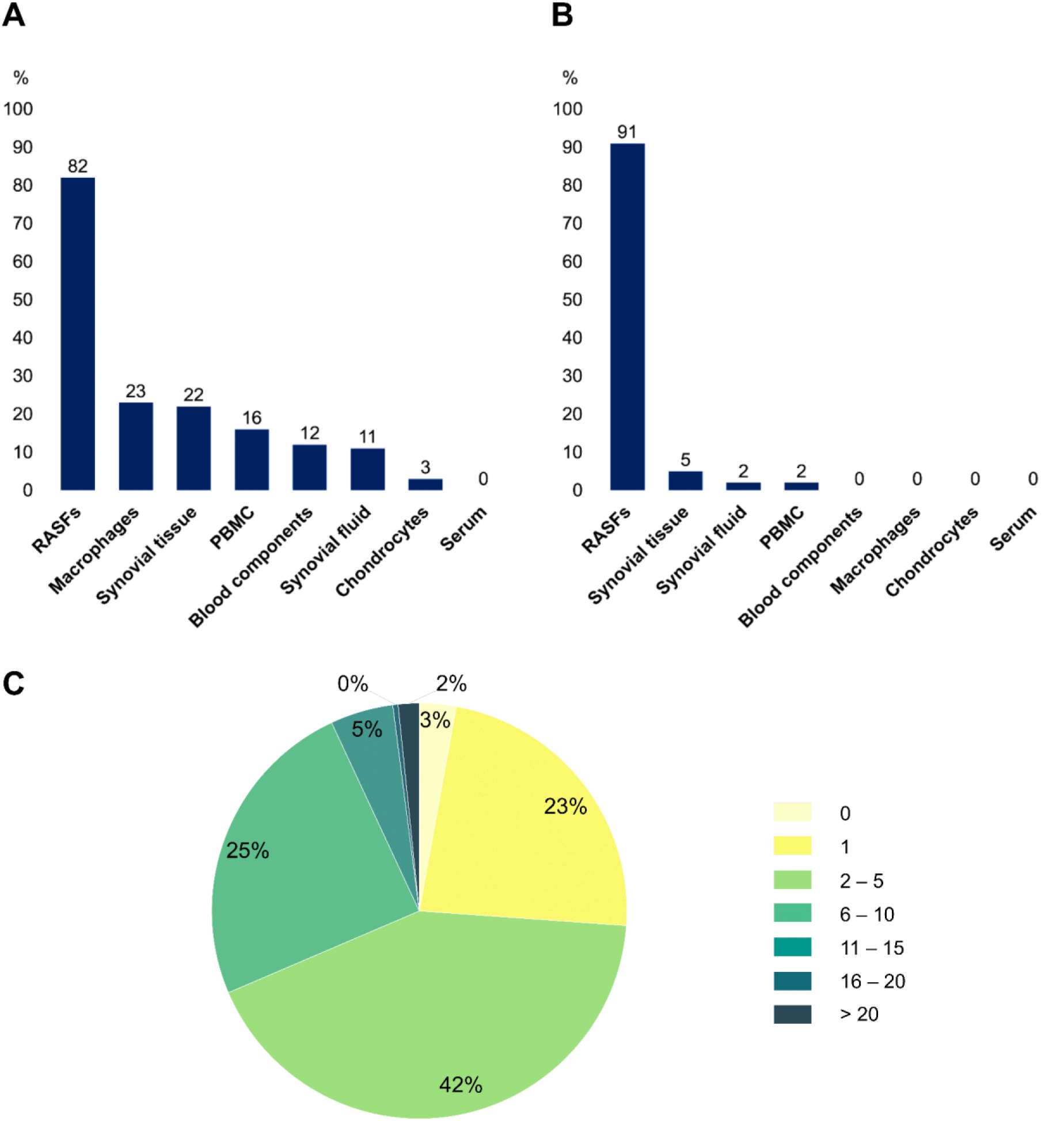
Statistical analysis of the RASF model. **(A)** Distribution of the RASF model cell-specificity. **(B)** Distribution of the RASF model cell-specificity when only exclusive components are considered. **(C)** Annotation score’s distribution among the RASF model’s components.

In addition, the RASF model’s annotation score enables high confidence in its knowledge base function. Indeed, 97% of the model’s components present more than one manually curated experimental evidence and 74% present more than 2 (Figure 2C). Various situations can be considered regarding the 3% of compounds for which no bibliographic reference is given. Many are simple molecules acting as products or reactants of well-known biological reactions whose expressions are rarely highlighted in disease-specific experimental studies (e.g., ATP, ADP, NADH, NADPH, H2O, O2, FADH, Ca2+). A further distinction is made regarding pathway intermediates. For instance, a specific pathway might be experimentally proven to be expressed in a disease-specific manner, but not all intermediates are necessarily studied. It is the case for several metabolic pathways that are found to be expressed in RASFs but experimental evidence is not available for every component.

To validate the RASF-specific behavior of the model, generic RASF model simulations were first compared to experimental scenarios (Table S2, appendix). Regarding this evaluation, 23 experimental scenarios were confirmed out of 30. The scenarios that were not reproducible were due to multiple reasons. First, information on the mechanism of certain interactions may be lacking in the literature, leading to a missing or incomplete representation in the map and the model. Also, the validation of some scenarios involved stoichiometric information that was not possible to reproduce with a strictly Boolean formalism. Finally, some generic scenarios were not validated since other pathways already activated/inactivated the said phenotype in the same initial conditions. Subsequent global model’s behavior evaluation under RASF-specific initial conditions (Table 2) confirmed RASFs’ experimental behavior. Indeed, aggressive phenotypes, i.e. bone erosion, cell chemotaxis, cell growth, hypoxia, inflammation, matrix degradation and osteoclastogenesis, are ON while apoptosis is OFF (Figure 3).

**Table 2.** RASF-specific initial conditions extracted from literature.

| Component | RASF-behavior | Source | Value |
| --- | --- | --- | --- |
| FASLG | Activated | Fukuda et al., 2021 | 1 |
| IL6 | Activated | Guerne et al., 1989 | 1 |
| IL18 | Activated | Gracie et al., 1999 | 1 |
| RANKL | Activated | Takayanagi et al., 2000 | 1 |
| MIR192 | Down-regulated | Li et al., 2017b | 0 |
| TNF | Activated | Lee et al., 2013 | 1 |
| FGF1 | Activated | Qu et al., 1995 | 1 |
| PDGFA | Activated | Rosengren et al., 2010 | 1 |
| WNT5A | Activated | Sen et al., 2001 | 1 |
| IL17A | Activated | Kotake et al., 1999 | 1 |
| TGFB1 | Activated | Cheon et al., 2002 | 1 |
| IKBA/NFKB1/RELA | Activated | Nejabatbakhsh Samimi et al., 2020 | 1 |
| SFRP5 | Down-regulated | Kwon et al., 2014 | 0 |
| GLC | Present | Garcia-Carbonell et al., 2016 | 1 |
| ATP | Present | Garcia-Carbonell et al., 2016 | 1 |

**Figure 3.**
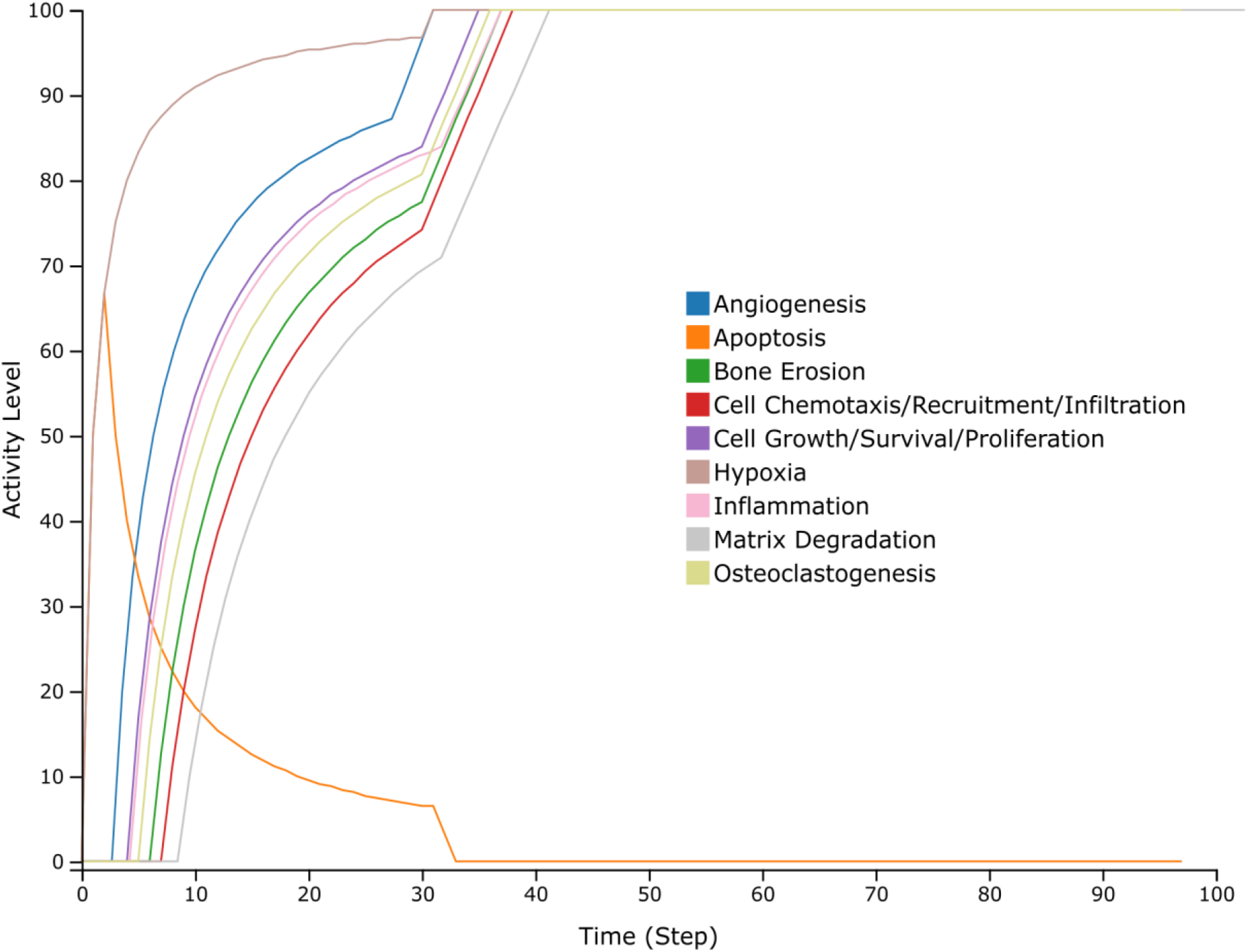
Simulation of the RASF model using RASF-specific initial conditions performed on the CellCollective platform. The different curves depict the state of the phenotype components.

### 3.2. Framework for hybrid modeling

The value propagation method was applied to the regulatory network using RASF-specific initial conditions extracted from the literature (Table 2). The results can be seen in Figure 4 in the form of a heatmap (Figure 4A) or mapped on the regulatory graph (Figure 4B). Of the 359 components present in the RASF model, 313 are fixed by value propagation. Among the 313 fixed values, 100 are fixed at 0 and 213 at 1. Evidently, the RASF-specific initial conditions exert an important control over the whole network.

**Figure 4.**
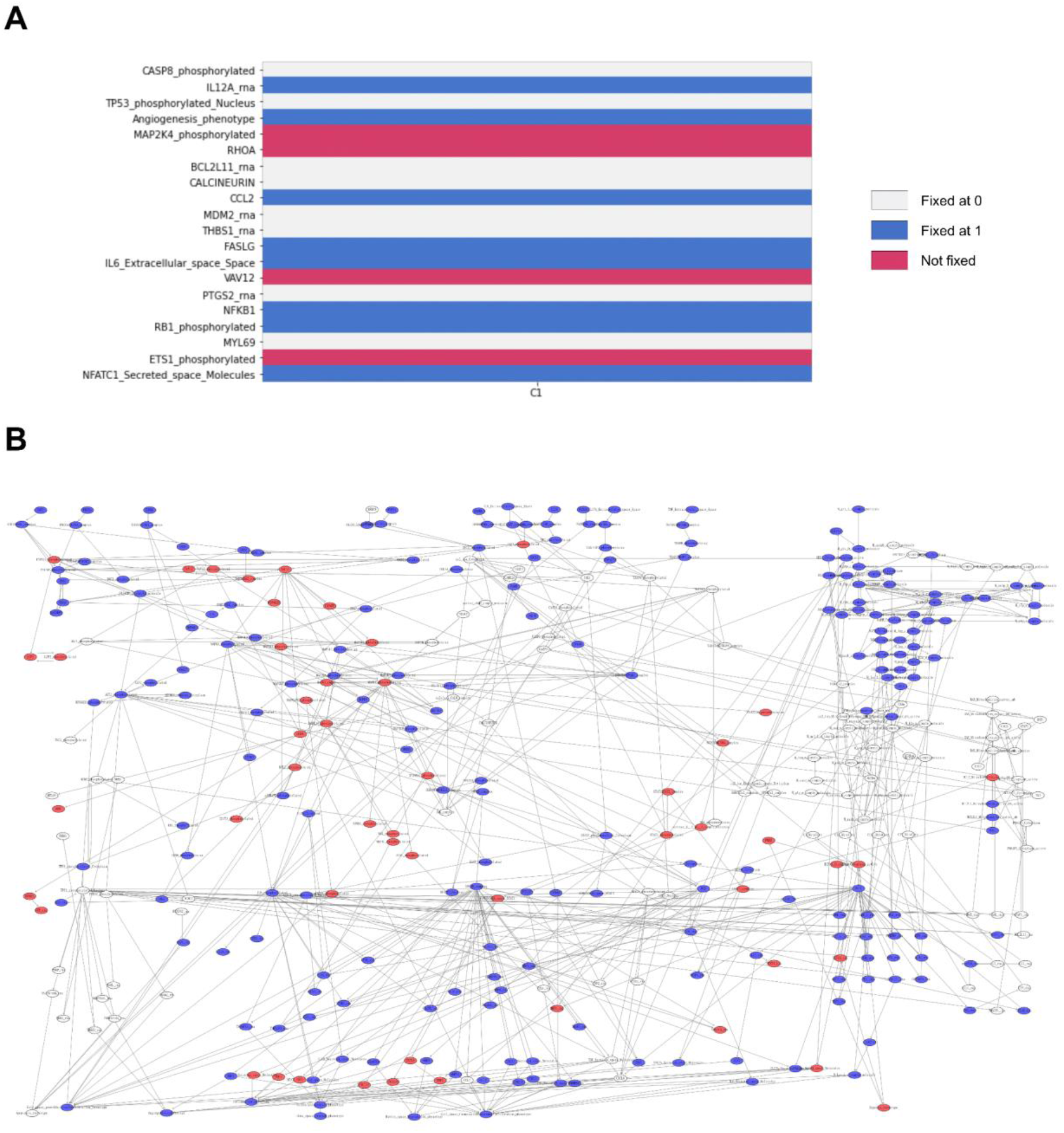
Results of value propagation performed on the RASF model using RASF-specific initial conditions. **(A)** Section of the heatmap of propagated values where each cell represents a component of the system. A white cell denotes a component fixed at value 0 in this condition. A blue cell denotes that it is fixed at 1. A red cell denotes components which are not fixed by value propagation in this condition. **(B)** Propagated values mapped on the RASF model’s regulatory network using the same color code.

The use of the results of value propagation as a new set of initial conditions enabled to decrease the complexity of the RASF model and obtain minimal trap-spaces which include the complete asymptotic behavior of the system (Table S3, appendix).

Using RASF-specific regulatory conditions, the maximal trap-spaces values for seven metabolic enzymes and 12 projected metabolites were equal to 0, constraining 52 metabolic reaction fluxes of the MitoCore model to 0. For instance, the maximal trap-space value relative to the metabolic enzyme AKGDm (2-Oxoglutarate Dehydrogenase) was equal to 0. Thus, the metabolic flux of the reaction it catalyzes (i.e., R_AKGDm) was constrained to 0 in MitoCore. Similarly, the maximal trap-space value associated with the metabolite fum_m (mitochondrial fumarate) was equal to 0. Consequently, the two metabolic reactions producing it (i.e., R_CII_MitoCore and R_FUMtmB_Mitocore) were constrained to 0 in MitoCore. Interestingly, several metabolic reactions’ fluxes (e.g., R_AKGDm, R_ICDHxm, R_CII_MitoCore, R_PDHm) were constrained to 0 in two distinct ways. Such reactions were constrained to 0 both by the extraction of metabolic constraints from the regulatory trap-spaces of the enzyme catalyzing the said reaction, and also from the metabolite produced by the said reaction. Such dually extracted constraints reflect the consistency of RASFs’ signaling and gene regulation’s impact on their metabolic pathways at different levels. For a detailed list of metabolic enzymes and metabolites with maximal regulatory trap-spaces values equal to 0 and the associated constrained reactions in MitoCore, refer to Tables S4 and S5 in the appendix.

### 3.3. Metabolic network analysis

A first FBA was carried to enable metabolic flux distribution’s assessment in a control situation. In addition, it allowed comparison with the second FBA, including metabolic constraints extracted from the RASF regulatory model. Results of both FBAs can be visualized in Figures 5A and 5B.

**Figure 5.**
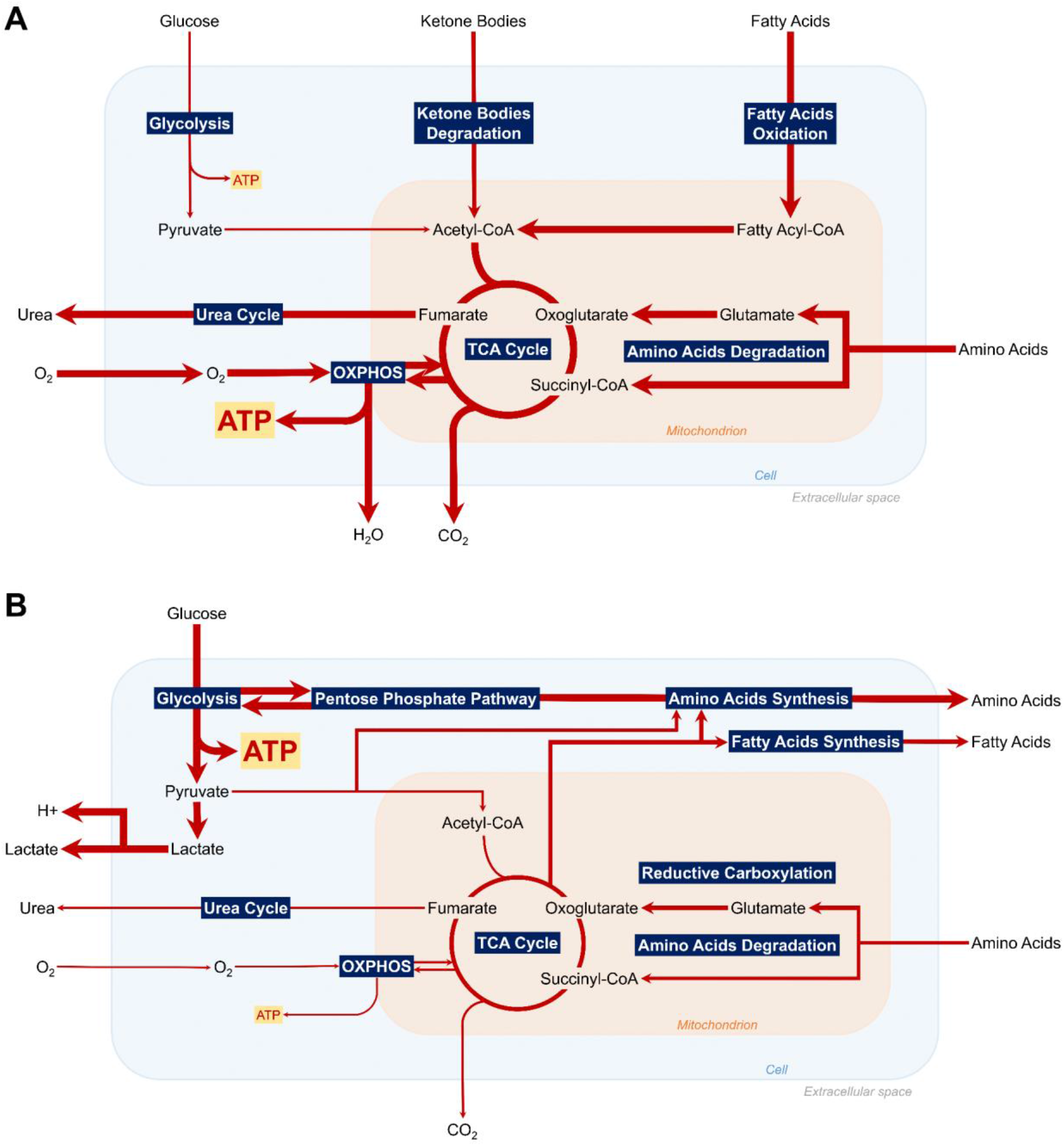
Summary of the major active pathways of central metabolism according to Flux Balance Analysis in control **(A)** and RASF-specific conditions **(B)** with maximum cellular ATP production as objective function.

As expected biologically, the optimal fluxes for maximum cellular ATP production in a control situation are TCA and OXPHOS fluxes. They are responsible for 96% of total ATP production. In a control situation, the main uptaken carbonated molecules are hexadecanoate (40.92% of total C-flux), glucose (29.48%), lactate (9.42%), and HCO3 (9.34%), primary energy sources for most cells. The main secreted carbonated molecule is CO2 (99.96%), the principal product of oxidative metabolism along with H2O. In RASF-specific regulatory conditions, the optimal fluxes for maximum cellular ATP production are glycolytic, accounting for 85% of cellular ATP production. Metabolic uptake and secretion fluxes are also affected. The main uptaken carbonated molecules are glucose (85.69% of total C-flux) and aspartate (9.77%). The main secreted carbonated molecules are lactate (86.12%) and alanine (8.33%). Details of the uptake and secretion C-fluxes results for both FBAs can be found in Tables S6-S9 in appendix.

In addition, a comparison of internal metabolic fluxes (Tables S10 and S11, appendix) in both situations illustrates increased glycolytic fluxes along with increased glucose uptake and lactate secretion in RASFs, accounting for a highly glycolytic metabolism. Low oxidative metabolism is demonstrated through decreased (almost null) TCA and OXPHOS fluxes and decreased secretion of OXPHOS by-products such as CO2 and H2O. A hypoxic environment is displayed with decreased O2 uptake and increased H^+^ secretion associated with environment acidity. Beyond metabolic pathways of ATP production, results denote reprogramming of several other metabolic pathways in RASFs. An increased amino-acids and fatty acids’ secretion is shown, potentially acting in RA as substrates for the production of energy through OXPHOS, biosynthesis intermediates, components of membrane phospholipids, or support for bone erosion and cartilage degradation. Increased reductive carboxylation is also identified, a novel glutamine metabolism pathway supporting the growth of tumor-like cells with mitochondrial defects. Further pathways including mitochondrial transport reactions, cardiolipin synthesis, or glycine cleavage, appear to be impacted, most likely indirectly as a result of metabolites’ redirection through up-regulated metabolic pathways.

To decipher the role of regulatory components in RASFs’ metabolic alterations, 14 FBAs were conducted following different sets of initial regulatory conditions (Table 3). As shown in Table 4, out of the 14 RASF-specific initial conditions variants, only condition 12 (C12) significantly impacted ATP production pathways. Indeed, when inhibiting Hypoxia-Inducible Factor 1 (HIF1) and keeping RASF-specific initial conditions for the remaining regulatory network’s inputs, glycolysis was dramatically decreased and OXPHOS explained the cellular ATP production. This situation, although extreme in its proportions due to the framework’s constraint extraction rules, is closer to a control situation. This finding suggests that targeting HIF1 could participate in restoring a healthy metabolic profile in RASFs. Moreover, it is coherent with recent experimental studies demonstrating that HIF1 knockdown reduces glycolytic metabolism in human synovial fibroblasts (Del Rey et al., 2017).

**Table 3.**
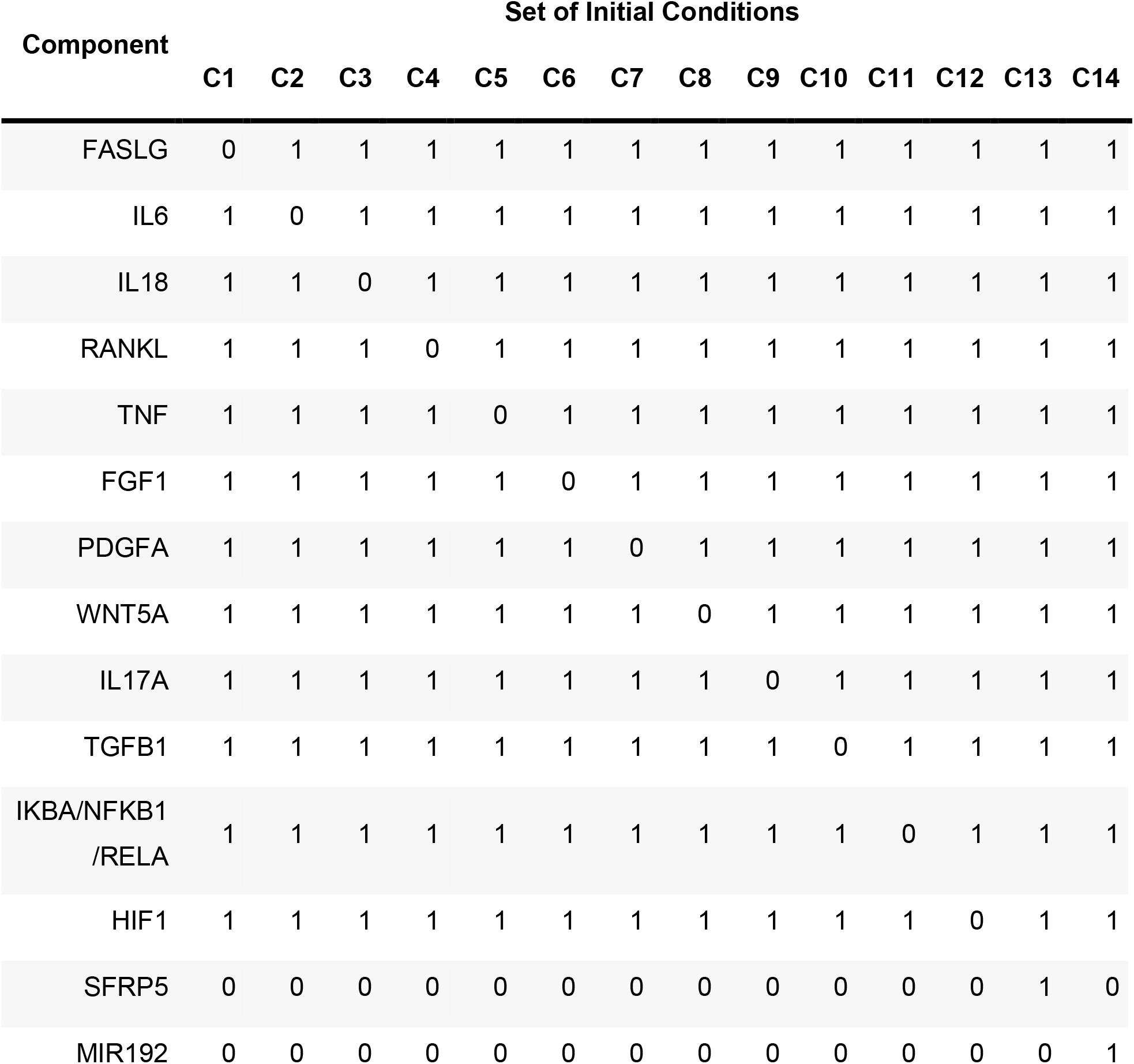
Summary of the 14 sets of RASF model’s initial conditions used for the regulatory knock-out/knock-in simulations. Each component’s value initially set to 1/0 in RASF-specific conditions was alternatively set to 0/1 while the others remained at RASF-specific values.

| Component | Set of Initial Conditions |  |  |  |  |  |  |  |  |  |  |  |  |  |
| --- | --- | --- | --- | --- | --- | --- | --- | --- | --- | --- | --- | --- | --- | --- |
|  | C1 | C2 | C3 | C4 | C5 | C6 | C7 | C8 | C9 | C10 | C11 | C12 | C13 | C14 |
| FASLG | 0 | 1 | 1 | 1 | 1 | 1 | 1 | 1 | 1 | 1 | 1 | 1 | 1 | 1 |
| IL6 | 1 | 0 | 1 | 1 | 1 | 1 | 1 | 1 | 1 | 1 | 1 | 1 | 1 | 1 |
| IL18 | 1 | 1 | 0 | 1 | 1 | 1 | 1 | 1 | 1 | 1 | 1 | 1 | 1 | 1 |
| RANKL | 1 | 1 | 1 | 0 | 1 | 1 | 1 | 1 | 1 | 1 | 1 | 1 | 1 | 1 |
| TNF | 1 | 1 | 1 | 1 | 0 | 1 | 1 | 1 | 1 | 1 | 1 | 1 | 1 | 1 |
| FGF1 | 1 | 1 | 1 | 1 | 1 | 0 | 1 | 1 | 1 | 1 | 1 | 1 | 1 | 1 |
| PDGFA | 1 | 1 | 1 | 1 | 1 | 1 | 0 | 1 | 1 | 1 | 1 | 1 | 1 | 1 |
| WNT5A | 1 | 1 | 1 | 1 | 1 | 1 | 1 | 0 | 1 | 1 | 1 | 1 | 1 | 1 |
| IL17A | 1 | 1 | 1 | 1 | 1 | 1 | 1 | 1 | 0 | 1 | 1 | 1 | 1 | 1 |
| TGFB1 | 1 | 1 | 1 | 1 | 1 | 1 | 1 | 1 | 1 | 0 | 1 | 1 | 1 | 1 |
| IKBA/NFKB1<br>/RELA | 1 | 1 | 1 | 1 | 1 | 1 | 1 | 1 | 1 | 1 | 0 | 1 | 1 | 1 |
| HIF1 | 1 | 1 | 1 | 1 | 1 | 1 | 1 | 1 | 1 | 1 | 1 | 0 | 1 | 1 |
| SFRP5 | 0 | 0 | 0 | 0 | 0 | 0 | 0 | 0 | 0 | 0 | 0 | 0 | 1 | 0 |
| MIR192 | 0 | 0 | 0 | 0 | 0 | 0 | 0 | 0 | 0 | 0 | 0 | 0 | 0 | 1 |

**Table 4.** Results of the regulatory initial condition’s knock-out/knock-in FBA simulations presented in terms of total cellular ATP production from glycolysis and OXPHOS.

| Set of Initial Conditions | Glycolysis (%) | OXPHOS (%) |
| --- | --- | --- |
| C1 | 85.05 | 14.95 |
| C2 | 85.05 | 14.95 |
| C3 | 85.05 | 14.95 |
| C4 | 85.05 | 14.95 |
| C5 | 85.05 | 14.95 |
| C6 | 85.05 | 14.95 |
| C7 | 85.05 | 14.95 |
| C8 | 85.05 | 14.95 |
| C9 | 85.05 | 14.95 |
| C10 | 85.05 | 14.95 |
| C11 | 85.05 | 14.95 |
| C12 | 0.00 | 100.00 |
| C13 | 85.05 | 14.95 |
| C14 | 85.05 | 14.95 |

## 4. Discussion

This work presents the construction of the first RASF-specific asynchronous Boolean model combining signaling, gene regulation, and metabolism. To obtain this model, the state-of-the-art RA-map V2 (Zerrouk et al., 2022) was used along with the map-to-model framework proposed by Aghamiri et al., 2020. The model was inferred based on RASF-specific pathways and components according to scientific literature. Additional biocuration ensured high cellular-specificity for RASFs. The model’s RASF-specific behavior was validated against biological scenarios extracted from the scientific literature and based on small-scale functional experiments. The integration of this Boolean model with a genomic-scale constraint-based metabolic model, MitoCore (Smith et al., 2017), allowed to assess the effects of RASFs’ regulatory outcomes on its central metabolic flux distribution.

To optimize the coupling of the two models, cell and disease-specific initial conditions were used to parametrize the regulatory model and decrease its complexity. The proposed framework only obtains constraints from metabolic compounds with a proven “inactive” asymptotic behavior, which allows it to automatically handle models with hundreds of components. This approach improves on previous attempts to couple Boolean models with metabolism, such as FlexFlux (Marmiesse et al., 2015b). In FlexFlux the discrete qualitative states of the Boolean regulatory network are translated into several user-defined continuous intervals, while in this framework only the metabolic reaction fluxes whose regulatory components have maximal trap-spaces equal to 0 are constrained. This choice is motivated by the difficulty of manually defining initial values and qualitative states to continuous intervals’ equivalence for every component of large regulatory models. In addition, this framework adopts the asynchronous update as less deterministic, and the identification of trap-spaces, using value propagation, to find states closer to the biological reality. This approach can facilitate the analysis of models with a higher number of inputs, however finding meaningful combinations of initial conditions can be challenging for models including less-studied entities. Moreover, regulatory models presenting less metabolic-associated maximal values of trap-spaces equal to 0, would obviously have fewer constraints applied to metabolic fluxes, decreasing the number of alterations of metabolic fluxes shown by FBA.

The results of the simulations of the hybrid RASF model for RASF-specific conditions revealed a highly glycolytic metabolism, along with a decreased oxidative metabolism for ATP production and confirmed a potential hypoxic environment in RASFs. The comparison with the simulation results for control FBA demonstrated clearly a metabolic reprogramming of RASFs. These results are consistent with recent experimental studies demonstrating a glycolytic switch in RASFs (de Oliveira et al., 2019; Fearon et al., 2018b).

Simulations of knock-outs and knock-ins of the regulatory initial conditions revealed HIF1 as a critical regulator of RASFs’ metabolic reprogramming. HIF1 is a master transcriptional factor involved in cellular and developmental response to hypoxia. Already identified in RA as a key effector of inflammation, angiogenesis, and cartilage destruction (Hua & Dias, 2016b; Gaber, 2005), HIF1 appears to be involved in RASFs’ metabolic alterations. In RASFs, HIF1 seems to promote glycolytic energy production by upregulating glucose transporters’ expression (i.e., GLUT1 and GLUT3) and transcription of enzymes responsible for intracellular glucose breakdown through glycolysis (e.g., Hexokinase, Phosphofructokinase-1, Aldolase), similarly to observations in CAFs (D. Zhang et al., 2015; Denko, 2008). In parallel, HIF1 might decrease ATP production through OXPHOS by transactivating genes responsible for O2 demand and mitochondrial activity, such as Pyruvate Dehydrogenase Kinase 1 or MAX Interactor 1. Considered together, these various properties enable HIF1 to enhance glycolysis’ rates as a crucial step of metabolic response to hypoxia (Denko, 2008).

RASFs’ metabolic alterations are generally attributed to their stressful microenvironment but could also be considered from the perspective of metabolite exchange between RASFs and other cells of the RA joint. Indeed, additional FBA results demonstrated several other altered metabolic pathways in RASFs, including fatty acids, amino acids and reductive carboxylation. The latter observations have not yet been experimentally studied in RASFs, but are similar to the ones made in Cancer-Associated Fibroblasts (CAFs). CAFs would undergo metabolic reprogramming to turn into “metabolic slaves”, generating energy-rich fuels and nutrients to feed neighboring cancer cells and help sustain their aggressive activity (Avagliano et al., 2018). Similarly, RASFs reside close to bio-energetically demanding cells (e.g., macrophages, dendritic cells, chondrocytes), experience a glycolytic switch and, according to these simulations, secrete high levels of energy-rich fuels and nutrients. The said nutrients are known to be involved in disease-associated behaviors, and some experimental evidence suggests intracellular metabolic exchanges between RASFs and neighboring cells (Bustamante et al., 2017b). Thus, a reverse Warburg relationship may occur between RASFs and RA joint cells. RASFs would undergo a metabolic switch and reprogram their metabolism to adapt to their hypoxic environment and provide crucial metabolic intermediates to neighboring cells to sustain their inflammatory activity. Further experimental studies are needed to decipher the intricate mechanisms of these metabolic exchanges, the precise beneficiaries of such metabolic intermediates, and the primary signal for this metabolic reprogramming.

## 5. Perspectives

In this approach, a “linear” view of events is adopted, with signaling, and gene regulation impacting metabolic pathways. However, the metabolic alterations due to HIF1’s regulatory action may initiate and contribute to a hypoxic feedback loop. To have a more realistic overview of RASFs’ dynamic behavior, further including the feedback of metabolism on gene regulation and signaling would be needed. An extension of this framework could be envisioned by adding a second timescale, as proposed in Thuillier et al., 2021, with the regulatory network getting updated once the metabolic network is at steady state. This extension would allow for a complete view of the metabolic reprogramming.

## 6. Conclusion

Hybrid modeling methods become increasingly important, especially in the era of systems biology, where biological phenomena are recognized as resulting from complex interactions at different layers. In RA, computational models able to span multiple biological processes could help decipher the intricate mechanisms at the origin of its pathogenesis. This work presents the first hybrid RASF model combining an RASF-specific asynchronous Boolean model with a global metabolic network. This hybrid RASF model is able to successfully reproduce the metabolic switch induced by signaling and gene regulation. Simulation results also enable further hypotheses on the potential reverse Warburg effect in RA. RASFs would reprogram their metabolism to produce more ATP, sustain their aggressive phenotype, and feed neighboring energetically demanding cells with fuels and nutrients. HIF1 was identified as the primary molecular switch driving RASFs’ metabolic reprogramming. Already recognized as an important target in the resolution of inflammation, targeting HIF1 might represent a promising path in the treatment of RA by addressing RASFs’ metabolic reprogramming.

## Availability

To facilitate reproducibility, we compiled all analysis in python notebooks and R scripts, in a step-by-step annotated manner. All data and code used to generate results are available on a GitLab repository at https://gitlab.com/genhotel/rasf-hybrid-model (accessed on July 13^th^ 2022).

## Notes

### Competing Interest Statement

The authors have declared no competing interest.

